# *Microbench*: Automated metadata management for systems biology benchmarking and reproducibility in Python

**DOI:** 10.1101/2021.09.14.460317

**Authors:** Alexander L. R. Lubbock, Carlos F. Lopez

**Affiliations:** Department of Biochemistry, Vanderbilt University, Nashville, Tennessee, United States of America; Vanderbilt-Ingram Cancer Center, Vanderbilt University, Nashville, Tennessee, United States of America; Department of Biomedical Informatics, Vanderbilt University Medical Center, Nashville, Tennessee, United States of America

## Abstract

**Motivation:** Computational systems biology analyses typically make use of multiple software and their dependencies, which often run across heterogeneous compute environments. This can introduce differences in performance and reproducibility. Capturing metadata (e.g. package versions, GPU model) currently requires repetitious code and is difficult to store centrally for analysis. Even where virtual environments and containers are used, updates over time mean that versioning metadata should still be captured within analysis pipelines to guarantee reproducibility.

**Results:** *Microbench* is a simple and extensible Python package to automate metadata capture to a file or Redis database. Captured metadata can include execution time, software package versions, environment variables, hardware information, Python version, and more, with plugins. We present three case studies demonstrating *Microbench* usage to benchmark code execution and examine environment metadata for reproducibility purposes.

**Availability:** Install from the Python Package Index using `pip install microbench`. Source code is available from https://github.com/alubbock/microbench.

## 1 Introduction

Analysis pipelines in computational biology frequently make use of multiple software dependencies and are often run across heterogenous compute environments (e.g. laptop, compute cluster, cloud). However, differences in software, hardware, and configuration can affect both reproducibility and performance. Standardized virtual environments using virtualenv (virtualenv.pypa.io), pipenv (pipenv.pypa.io), or Anaconda (anaconda.com) are an important first step towards reproducibility. Containerization frameworks like Docker (Boettiger, 2015) further help by standardizing system libraries and adding portability, but require time to setup and compile, and so are not always used during development. Even with virtual environments and containers, it remains important to track versions as the environments and containers themselves are updated over time. In addition, Python *requirements.txt* files often only specify the direct dependencies of the code, and not downstream dependencies which may change if an environment or container is re-built at a later date. Continuous integration (CI) (Meyer, 2014) can help to identify these issues, but reproducibility issues can also occur due to differences in hardware (e.g. GPU model) and environment (e.g. environment variables). In addition, comprehensive CI is not always feasible with large -omics datasets. Capturing the full environment at runtime would help to identify the cause of reproducibility issues, should they be discovered later.

Here, we introduce *microbench*, an open source Python package which automates metadata capture and execution timing of annotated functions for performance bench-marking and to improve reproducibility. For example, *microbench* can capture Python package versions, environment variables, host hardware specifications, Python version, and function arguments. It can capture line-by-line execution times and telemetry – CPU and RAM utilization recorded periodically during function execution. Results are logged to a file or a central Redis database (redis.io), and can be analyzed using *pandas* DataFrames (pandas.pydata.org). *Microbench* can be extended with plugins to capture additional metadata types. *Microbench* lowers the administrative burden of capturing metadata for benchmarking and reproducibility. We believe it will significantly help computational biologists working in Python who faces these challenges.

## 2 Results

### Implementation

*Microbench* is written in pure Python for cross-platform compatibility (tested on Windows 10, Mac, Linux), and has no default runtime dependencies outside of the Python standard library. However, the *line_profiler* package (pypi.org/project/line-profiler) is required for line-by-line benchmarking, and *pandas* is recommended to examine and analyze results. The package is designed to be easy to use and is extensible with new metadata capture capabilities.

### Installation and usage

*Microbench* is installed using `pip install microbench` (Supplementary Text S1). Source code is available on GitHub (https://github.com/alubbock/microbench) under the MIT license. A minimal usage example is given in Supplementary Text S2; an extended usage example is given in Supplementary Text S3. Briefly, a benchmark suite is first specified, which determines what metadata are captured. A Python decorator is then used to mark functions for benchmarking/metadata capture. Results are saved to a JSON file or Redis instance for later analysis (Figure 1). The results format is described in Supplementary Text S4.

**Figure 1:**
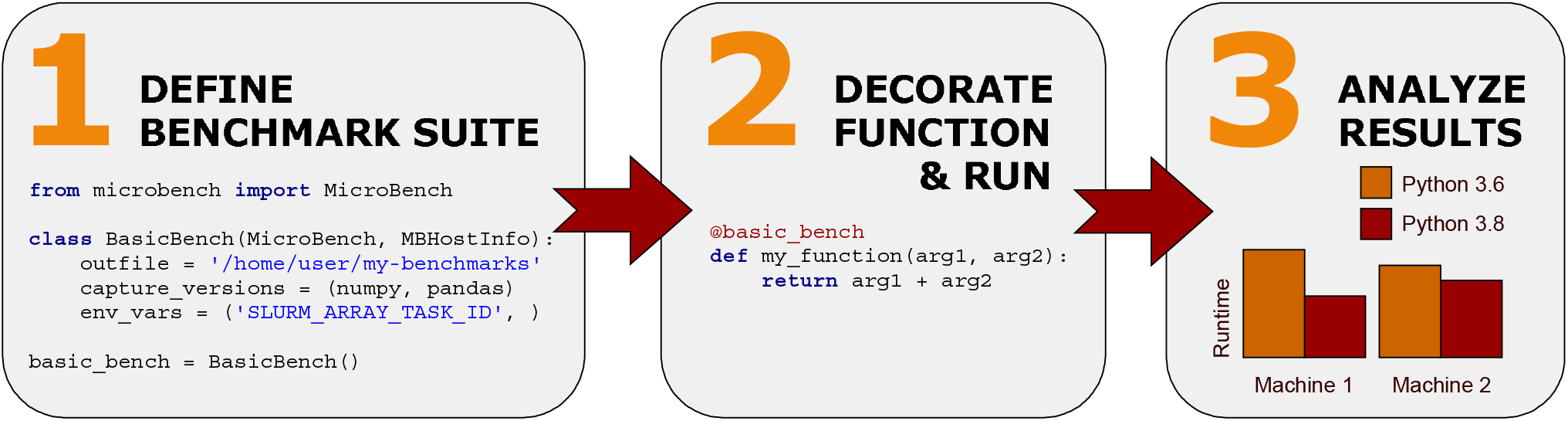
*Microbench* usage. The user defines the set of metadata to capture, then decorates the target function and runs it, then analyzes the results. Captured metadata can be used to check issues in case of a reproducibility anomaly, or for performance benchmarking.

### Metadata capture

*Microbench* can capture a wide variety of metadata. By default, start timestamp, end timestamp, and function name are captured. Additional metadata are determined by the user in the form of mixins – small Python classes which are included when the benchmark suite is constructed. A list of available mixins is shown in Supplementary Table 1. *Microbench* can be extended with custom mixins to capture additional or bespoke metadata (Supplementary Text S5).

### Redis support

*Microbench* typically appends data to a file, which works well for single machine use cases or where a shared filesystem can be used. However, when using the cloud or distributed computing platforms, collecting metadata from multiple files is inefficient. *Microbench* includes support for Redis – an unstructured, in-memory database. If Redis is run on an internet-accessible server, metadata can be deposited from any internet-connected device. *Microbench* Redis support requires the redis-py package (pypi.org/project/redis) (Supplementary Text S6).

### NVIDIA GPU support

Attributes relevant to NVIDIA GPUs can be captured using a metadata collector for NVIDIA’s nvidia-smi utility (developer.nvidia.com/nvidia-system-management-interface), including GPU model number and GPU memory capacity (Supplementary Text S7).

### Line profiler support

The ability to capture line-by-line execution times is useful in several situations, but particularly in algorithm development, where line profiling can help to identify bottlenecks for targeted improvements. *Microbench* can capture these data by integration with the line_profiler package (Supplementary Text S8).

### Telemetry support

*Microbench* can capture function telemetry – metrics such as CPU and RAM utilization, captured at specified intervals by a background thread. This feature can help diagnose bottlenecks and resource intensive sections of code, particularly in conjunction with the line_profiler (Supplementary Text S9).

### Case study: biochemical model simulation

We include an application of *microbench* to a biochemical model simulation using PySB (Lopez et al., 2013) within a Jupyter Notebook (Perkel, 2018), where we capture package versions from an Anaconda environment to check for reproducibility (Supplementary File S1, example 1). This example also uses telemetry analysis to examine CPU and RAM utilization over time during function execution.

### Case study: NumPy return value differs by version

Software library updates can result in differences or errors– which may or may not be documented in those libraries’ documentation or spotted by the end user. We present a simple example case study where a computation gives a different result using NumPy 1.15 compared to NumPy 1.20 (https://numpy.org/doc/stable/release/1.20.0-notes.html#np-linspace-on-integers-now-uses-floor). Capturing version metadata can help to identify such issues retrospectively, when required (Supplementary File S1, example 2).

### Case study: opportunistic benchmarking using SLURM

SLURM (Yoo et al., 2003) is a cluster management system which can be used to execute computational jobs in parallel. Large clusters are often subject to rolling or phased hardware refreshes, leading to a heterogeneous compute environment. *Microbench* can capture hardware specifications, and subsequently the runtime of an algorithm can be compared by feature (e.g. max. CPU frequency, total RAM) for benchmarking or specifying requirements (Supplementary File S1, example 3). Bench-marking can be “opportunistic” in the sense that relevant metadata can be captured with little additional effort, in case of later need.

## 3 Discussion

*Microbench* is designed to provide a simple and unobtrusive way to benchmark Python functions and capture key metadata for reproducibility. It is particularly useful in heterogenous compute environments and ephemeral cloud computing environments, where one cannot easily go back and examine the exact instance after analysis completion. *Microbench* has no runtime dependencies outside the Python standard library, making it easy to deploy across heterogeneous compute environments. By making metadata capture simple to add to existing Python code, we hope users will use it routinely, allowing for retrospective debugging and benchmarking should an issue arise later.

