## Supplementary Material for "*Microbench*: Automated metadata management for systems biology benchmarking and reproducibility in Python"

### Supplementary Information

#### Supplementary Text S1: Requirements and Installation

Microbench by default has no dependencies outside of the Python standard library, although pandas is recommended to examine results. However, some mixins (extensions) have specific requirements:

- The line_profiler package needs to be installed for line-by-line code benchmarking.
- `MBInstalledPackages` requires `setuptools`, which is not a part of the standard library, but is usually available.
- The CPU cores, total RAM, and telemetry extensions require psutil.
- The NVIDIA GPU plugin requires the nvidia-smi utility, which usually ships with the NVIDIA graphics card drivers. It needs to be on your `PATH`.

To install using `pip`:

```
pip install microbench
```

#### Supplementary Text S2: Minimal usage example

Microbench is designed for benchmarking Python functions. These examples will assume you have already defined a Python function `myfunction` that you wish to benchmark:

```
def myfunction(arg1, arg2, ...):
    ...
```

First, create a benchmark suite, which specifies the configuration and information to capture.

Here's a minimal, complete example:

```
from microbench import MicroBench

basic_bench = MicroBench()
```

To attach the benchmark to your function, simply use `basic_bench` as a decorator, like this:

```
@basic_bench
def myfunction(arg1, arg2, ...):
```

```
    ...
```

That's it! When `myfunction()` is called, metadata will be captured into a `io.StringIO()` buffer, which can be read as follows (using the `pandas` library):

```
import pandas as pd
results = pd.read_json(basic_bench.outfile.getvalue(), lines=True)
```

The above example captures the fields `start_time`, `finish_time` and `function_name`. Microbench can capture many other types of metadata from the environment, resource usage, and hardware, which are covered below.

### Supplementary Text S3: Extended usage example

Here's a more complete example using mixins (the `MB` prefixed class names) to extend functionality. Note that keyword arguments can be supplied to the constructor (in this case `some_info=123`) to specify additional information to capture. This example also specifies the `outfile` option, which appends metadata to a file on disk.

```
from microbench import *
import numpy, pandas

class MyBench(MicroBench, MBFunctionCall, MBPythonVersion, MBHostInfo):
    outfile = '/home/user/my-benchmarks'
    capture_versions = (numpy, pandas)  # Or use
MBGlobalPackages/MBInstalledPackages
    env_vars = ('SLURM_ARRAY_TASK_ID', )

benchmark = MyBench(some_info=123)
```

The `env_vars` option from the example above specifies a list of environment variables to capture as `env_<variable name>`. In this example, the slurm array task ID will be stored as `env_SLURM_ARRAY_TASK_ID`. Where the environment variable is not set, the value will be `null`.

To capture package versions, you can either specify them individually (as above), or you can capture the versions of every package in the global environment. In the following example, we would capture the versions of `microbench`, `numpy`, and `pandas` automatically.

```
from microbench import *
import numpy, pandas

class Bench2(MicroBench, MBGlobalPackages):
    outfile = '/home/user/bench2'

bench2 = Bench2()
```

If you want to go even further, and capture the version of every package available for import, there's a mixin for that:

```
from microbench import *

class Bench3(MicroBench, MBInstalledPackages):
```

```
    pass

bench3 = Bench3()
```

### Supplementary Text S4: Examine microbench results files

Each result is a JSON object. When using the `outfile` option, a JSON object for each `@benchmark` call is stored on a separate line in the file. The output from the minimal example above for a single run will look similar to the following:

```
{"start_time": "2018-08-06T10:28:24.806493", "finish_time": "2018-08-06T10:28:24.867456", "function_name": "my_function"}
```

The simplest way to examine results in detail is to load them into a pandas dataframe:

```
import pandas
results = pandas.read_json('/home/user/my-benchmarks', lines=True)
```

Pandas has powerful data manipulation capabilities. For example, to calculate the average runtime by Python version:

```
# Calculate runtime for each run
results['runtime'] = results['finish_time'] - results['start_time']

# Average runtime by Python version
results.groupby('python_version')['runtime'].mean()
```

Many more advanced operations are available. The pandas tutorial is recommended.

### Supplementary Text S5: Extending microbench

Microbench includes a few mixins for basic functionality as described in the extended example, above.

You can also add functions to your benchmark suite to capture extra information at runtime. These functions must be prefixed with `capture_` for them to run automatically before the function starts. They take a single argument, `bm_data`, a dictionary to be extended with extra data. Care should be taken to avoid overwriting existing key names.

Here's an example to capture the machine type (`i386`, `x86_64` etc.):

```
from microbench import MicroBench
import platform

class Bench(MicroBench):
    outfile = '/home/user/my-benchmarks'

    def capture_machine_platform(self, bm_data):
        bm_data['platform'] = platform.machine()

benchmark = Bench()
```

### Supplementary Text S6: Redis support

By default, microbench appends output to a file, but output can be directed elsewhere, e.g. redis - an in-memory, networked data source. This option is useful when a shared filesystem is not available.

Redis support requires redis-py.

To use this feature, inherit from `MicroBenchRedis` instead of `MicroBench`, and specify the redis connection and key name as in the following example:

```
from microbench import MicroBenchRedis

class RedisBench(MicroBenchRedis):
    # redis_connection contains arguments for redis.StrictClient()
    redis_connection = {'host': 'localhost', 'port': 6379}
    redis_key = 'microbench:mykey'

benchmark = RedisBench()
```

To retrieve results, the `redis` package can be used directly:

```
import redis
import pandas

# Establish the connection to redis
rconn = redis.StrictRedis(host=..., port=...)

# Read the redis data from 'myrediskey' into a list of byte arrays
redis_data = redis.lrange('myrediskey', 0, -1)

# Convert the list into a single string
json_data = '\n'.join(r.decode('utf8') for r in redis_data)

# Read the string into a pandas dataframe
results = pandas.read_json(json_data, lines=True)
```

### Supplementary Text S7: NVIDIA GPU support

Attributes about NVIDIA GPUs can be captured using the `MBNvidiaSmi` plugin. This requires the `nvidia-smi` utility to be available in the current `PATH`.

By default, the `gpu_name` (model number) and `memory.total` attributes are captured. Extra attributes can be specified using the class or object-level variable `nvidia_attributes`. To see which attributes are available, run `nvidia-smi --help-query-gpu`.

By default, all installed GPUs will be polled. To limit to a specific GPU, specify the `nvidia_gpus` attribute as a tuple of GPU IDs, which can be zero-based GPU indexes (can change between reboots, not recommended), GPU UUIDs, or PCI bus IDs. You can find out GPU UUIDs by running `nvidia-smi -L`.

Here's an example specifying the optional `nvidia_attributes` and `nvidia_gpus` fields:

```
from microbench import MicroBench, MBNvidiaSmi

class GpuBench(MicroBench, MBNvidiaSmi):
    outfile = '/home/user/gpu-benchmarks'
    nvidia_attributes = ('gpu_name', 'memory.total', 'pcie.link.width.max')
    nvidia_gpus = (0, )  # Usually better to specify GPU UUIDs here instead

gpu_bench = GpuBench()
```

### Supplementary Text S8: Line profiler support

Microbench also has support for line_profiler, which shows the execution time of each line of Python code. Note that this will slow down your code, so only use it if needed, but it's useful for discovering bottlenecks within a function. Requires the `line_profiler` package to be installed (e.g. `pip install line_profiler`).

```
from microbench import MicroBench, MBLineProfiler
import pandas

# Create our benchmark suite using the MBLineProfiler mixin
class LineProfilerBench(MicroBench, MBLineProfiler):
    pass

lpbench = LineProfilerBench()

# Decorate our function with the benchmark suite
@lpbench
def my_function():
    """ Inefficient function for line profiler """
    acc = 0
    for i in range(1000000):
        acc += i

    return acc

# Call the function as normal
my_function()

# Read the results into a Pandas DataFrame
results = pandas.read_json(lpbench.outfile.getvalue(), lines=True)

# Get the line profiler report as an object
lp = MBLineProfiler.decode_line_profile(results['line_profiler'][0])

# Print the line profiler report
MBLineProfiler.print_line_profile(results['line_profiler'][0])
```

The last line of the previous example will print the line profiler report, showing the execution time of each line of code. Example:

```
Timer unit: 1e-06 s

Total time: 0.476723 s
File: /home/user/my_test.py
Function: my_function at line 12

Line #      Hits         Time  Per Hit   % Time  Line Contents
==============================================================
```

```
    12                                                     @lpbench
    13                                                     def my_function():
    14                                                         """ Inefficient
function for line profiler """
    15         1          2.0      2.0      0.0          acc = 0
    16   1000001     217874.0      0.2     45.7          for i in
range(1000000):
    17   1000000     258846.0      0.3     54.3              acc += i
    18
    19         1          1.0      1.0      0.0          return acc
```

### Supplementary Text S9: Telemetry support

We use the term "telemetry" to refer to metadata which is captured periodically during the execution of a function by a thread which runs in parallel. For example, this may be useful to see how memory usage changes over time.

Telemetry support requires the `psutil` library.

Microbench implements the thread automatically; the end user only need define what metadata are captured and return the result as a dictionary. The default telemetry collection interval is every 60 seconds, which can be customized if needed.

A minimal example to capture memory usage every 90 seconds is shown below:

```
from microbench import MicroBench

class TelemBench(MicroBench):
    @staticmethod
    def telemetry(process):
        return process.memory_full_info()._asdict()

telem_bench = TelemBench()
```

### Supplementary Table S1: List of Microbench mixins

**Supplementary Table S1.** List of *Microbench* mixins (metadata capture extensions)

| Mixin | Fields captured |
|---|---|
| *(default)* | `start_time`<br>`finish_time`<br>`function_name` |
| MBGlobalPackages | `package_versions`, with entry for every package in the global environment |
| MBInstalledPackages | `package_versions`, with entry for every package available for import |
| MBCondaPackages | `conda_versions`, with entry for every conda package in the environment |
| MBFunctionCall | `args` (positional arguments)<br>`kwargs` (keyword arguments) |
| MBReturnValue | Wrapped function's return value |
| MBPythonVersion | `python_version` (e.g. 3.6.0)<br>`python_executable` |

| | |
|---|---|
| | (e.g. `/usr/bin/python`, which should indicate any active virtual environment) |
| MBHostInfo | `hostname`<br>`operating_system` |
| MBHostCpuCores | `cpu_cores_logical`<br>(number of cores, requires `psutil`) |
| MBHostRamTotal | `ram_total`<br>(total RAM in bytes, requires `psutil`) |
| MBNvidiaSmi | Various NVIDIA GPU fields, see Supplementary Text 7 |
| MBLineProfiler | `line_profiler` containing line-by-line profile, see Supplementary Text S8 |
