## Supplementary File S1 for "*Microbench*: Automated metadata management for systems biology benchmarking and reproducibility in Python": python-package.yml

```
# This workflow will install Python dependencies, run tests and lint with a variety of Python versions
# For more information see: https://help.github.com/actions/language-and-framework-guides/using-python-with-github-actions

name: Python package

on:
  push:
    branches:
      - main
  pull_request:
    branches:
      - main

jobs:
  build:
    runs-on: ubuntu-latest
    defaults:
      run:
        shell: bash -l {0}

    steps:
    - uses: actions/checkout@v2
    - name: Install miniconda environment
      uses: conda-incubator/setup-miniconda@v2
      with:
        activate-environment: microbench-examples
        environment-file: environment.yml
        python-version: 3.9
    - name: conda info
      run: conda info
    - name: Test with pytest
      run: |
        pytest
```
