## Supplementary File S1 for "*Microbench*: Automated metadata management for systems biology benchmarking and reproducibility in Python": environment.yml

```
name: microbench-examples
channels:
  - alubbock
  - defaults
dependencies:
  - bionetgen=2.5.1
  - blas=1.0
  - ca-certificates=2021.5.25
  - certifi=2021.5.30
  - gmp=6.2.1
  - gmpy2=2.0.8
  - intel-openmp=2021.2.0
  - libffi=3.3
  - mkl=2021.2.0
  - mkl-service=2.3.0
  - mkl_fft=1.3.0
  - mkl_random=1.2.1
  - mpc=1.1.0
  - mpfr=4.0.2
  - mpmath=1.2.1
  - ncurses=6.2
  - networkx=2.5.1
  - nfsim=1.12.1
  - numpy-base=1.20.2
  - openssl=1.1.1k
  - pip=21.1.3
  - pysb=1.13.0
  - python=3.9.5
  - python-dateutil=2.8.1
  - readline=8.1
  - setuptools=52.0.0
  - six=1.16.0
  - sqlite=3.36.0
  - sympy=1.8
  - tk=8.6.10
  - tzdata=2021a
  - wheel=0.36.2
  - xz=5.2.5
  - zlib=1.2.11
  - pip:
    - appnope==0.1.2
    - attrs==21.2.0
    - backcall==0.2.0
    - coverage==5.5
    - cycler==0.10.0
    - debugpy==1.3.0
    - decorator==5.0.9
    - iniconfig==1.1.1
    - ipykernel==6.0.0
    - ipython==7.25.0
    - ipython-genutils==0.2.0
    - jedi==0.18.0
    - jsonschema==3.2.0
    - jupyter-client==6.1.12
    - jupyter-core==4.7.1
    - kiwisolver==1.3.1
    - matplotlib==3.4.2
    - matplotlib-inline==0.1.2
    - microbench==0.7
    - nbformat==5.1.3
    - nbval==0.9.6
    - numpy==1.21.0
    - packaging==20.9
    - pandas==1.2.5
    - parso==0.8.2
    - pexpect==4.8.0
    - pickleshare==0.7.5
    - pillow==8.2.0
    - pluggy==0.13.1
    - prompt-toolkit==3.0.19
    - psutil==5.8.0
    - ptyprocess==0.7.0
    - py==1.10.0
    - pygments==2.9.0
    - pyparsing==2.4.7
    - pyrsistent==0.18.0
    - pytest==6.2.4
    - pytz==2021.1
    - pyzmq==22.1.0
    - scipy==1.7.0
    - seaborn==0.11.1
    - toml==0.10.2
    - tornado==6.1
    - traitlets==5.0.5
    - wcwidth==0.2.5
```
