## Supplementary File S1 for "*Microbench*: Automated metadata management for systems biology benchmarking and reproducibility in Python": microbench-pysb.ipynb

```
# %% [markdown]
# Analyzing benchmarks of EARM 1.0 PySB model with microbench

# %% [markdown]
## Background

[PySB](https://pysb.org) is a biochemical modeling framework for Python. Here, we analyze data from running simulations of a model ([Extrinsic apoptosis reaction model version 1.0](https://github.com/pysb/pysb/blob/master/pysb/examples/run_earm_1_0.py)) using an ODE integrator from the SciPy library. We examine the runtime across two different functions, and check that the environment (packages and their versions) are consistent across all runs, for reproducibility purposes.

# %% [markdown]
## Load the data

# %% [code]
%matplotlib inline

# %% [code]
import pandas
import seaborn
import numpy

# %% [code]
# Show up to 200 rows in a dataframe
pandas.set_option('display.max_rows', 200)

# %% [code]
results = pandas.read_json('microbench-pysb.json', lines=True)

# %% [markdown]
## Calculate mean runtime by function

# %% [code]
# Calculate runtime for each run
results['runtime'] = results['finish_time'] - results['start_time']

# Average runtime by function name
print(results.groupby('function_name')['runtime'].mean(numeric_only=False))

# %% [markdown]
## Show mean runtime as a boxplot with whiskers

# %% [code]
seaborn.boxplot(x=results['function_name'], y=results['runtime'].dt.total_seconds())

# %% [markdown]
## Check conda environment consistency

We can quickly check if the conda environment was consistent across runs. We convert the conda_versions data (stored as dictionaries) to strings, and check that they're the same for every run:

# %% [code]
len(results['conda_versions'].astype('str').unique()) == 1

# %% [markdown]
We can also convert the package versions from a single run into a pandas dataframe:

# %% [code]
pandas.DataFrame.from_dict(results['conda_versions'][0], orient='index', columns=['Version'])

# %% [markdown]
Microbench provides a function, `envdiff`, to compare two environments together in a Jupyter Notebook, with differences highlighted in red (scroll to bottom in the following example). 

# %% [code]
from microbench import envdiff
envdiff(results.iloc[0], results.iloc[1])

# %% [markdown]
## Telemetry analysis
The function runs above contain a few telemetry points. Let's see how CPU and RAM usage changed over time in the first run.

# %% [code]
run0_telem = pandas.DataFrame.from_dict(results.iloc[0].telemetry)
# Convert timestamps from str to datetime64
run0_telem['timestamp'] = pandas.to_datetime(run0_telem['timestamp'])
# Show the dataframe
run0_telem

# %% [code]
# Calculate elapsed time for each entry
run0_telem['time_elapsed'] = run0_telem['timestamp'] - results.iloc[0]['start_time']

# %% [code]
# Show system CPU percent utilization over time
seaborn.lineplot(data=run0_telem, y='sys_cpu_percent', x='time_elapsed')

# %% [code]
# Show resident set size (RSS) memory usage over time
run0_telem['process_rss_mem'] = run0_telem['process_mem'].apply(lambda x: x[0])
seaborn.lineplot(data=run0_telem, y='process_rss_mem', x='time_elapsed')

# %% [markdown]
## Discussion
The above examples show how multiple functions can be benchmarked using microbench, and how different types of metadata can be captured. We also showed how to display telemetry data on a contrived example, but the same principles will apply to telemetry from longer-running functions

Microbench aims to make metadata capture easy, so that it can be collected routinely and used later if required for debugging, identifying reproducibility issues, and benchmarking.
```
