## Supplementary File S1 for "*Microbench*: Automated metadata management for systems biology benchmarking and reproducibility in Python": microbench_pysb.py

```
"""Reproduce figures 4A and 4B from the EARM 1.0 publication (Albeck et
al. 2008)."""

from __future__ import print_function
from pysb.simulator import ScipyOdeSimulator
import matplotlib.pyplot as plt
from numpy import *

from pysb.examples.earm_1_0 import model

# Benchmarking code
from microbench import MicroBench, MBCondaPackages, MBHostInfo
import pandas
import psutil


class EarmBench(MicroBench, MBCondaPackages, MBHostInfo):
    outfile = 'microbench_pysb.json'
    telemetry_interval = 2

    @staticmethod
    def telemetry(process):
        with process.oneshot():
            return {
                'process_cpu_percent': process.cpu_percent(interval=0.1),
                'process_mem': process.memory_full_info(),
                'sys_swap': psutil.swap_memory(),
                'sys_mem': psutil.virtual_memory(),
                'cpu_freq_current': psutil.cpu_freq().current,
                'sys_cpu_percent': psutil.cpu_percent(interval=0.1)
            }


earmbench = EarmBench()
# End benchmarking code

# saturating level of ligand (corresponding to ~1000 ng/ml SuperKiller TRAIL)
Lsat = 6E4

# relationship of ligand concentration in the model (in # molecules/cell)
# to actual TRAIL concentration (in ng/ml)
Lfactor = model.parameters['L_0'].value / 50

L_0_baseline = model.parameters['L_0'].value

sim = ScipyOdeSimulator(model)


@earmbench
def fig_4a():
    print("Simulating model for figure 4A...")
    t = linspace(0, 20*3600, 20*60+1)  # 20 hours, in seconds, 1 min sampling
    dt = t[1] - t[0]

    Ls_exp = Lsat / array([1, 4, 20, 100, 500])
    Td_exp = [144.2, 178.7, 236,   362.5, 656.5]
    Td_std = [32.5,   32.2,  36.4,  78.6, 171.6]
    Ts_exp = [21.6,   23.8,  27.2,  22.0,  19.0]
    Ts_std = [9.5,     9.5,  12.9,   7.7,  10.5]

    CVenv = 0.2
    # num steps was originally 40, but 15 is plenty smooth enough for screen display
    Ls = floor(logspace(1,5,15))

    fs = empty_like(Ls)
    Ts = empty_like(Ls)
    Td = empty_like(Ls)
    print("Scanning over %d values of L_0" % len(Ls))
    for i, L_0 in enumerate(Ls):
        print("  L_0 = %g" % L_0)
        x = sim.run(tspan=t, param_values={"L_0": L_0}).all

        fs[i] = (x['PARP_unbound'][0] - x['PARP_unbound'][-1]) / x['PARP_unbound'][0]
        dP = 60 * (x['PARP_unbound'][:-1] - x['PARP_unbound'][1:]) / (dt * x['PARP_unbound'][0])  # in minutes
        ttn = argmax(dP)
        dPmax = dP[ttn]
        Ts[i] = 1 / dPmax  # minutes
        Td[i] = t[ttn] / 60  # minutes

    plt.figure("Figure 4A")
    plt.plot(Ls/Lfactor, Td, 'g-', Ls/Lfactor, (1-CVenv)*Td, 'g--', Ls/Lfactor,
          (1+CVenv)*Td, 'g--')
    plt.errorbar(Ls_exp/Lfactor, Td_exp, Td_std, None, 'go', capsize=0),
    plt.ylabel('Td (min)'),
    plt.xlabel('TRAIL (ng/ml)'),
    a = plt.gca()
    a.set_xscale('log')
    a.set_xlim((min(Ls) / Lfactor, max(Ls) / Lfactor))
    a.set_ylim((0, 1000))

    model.parameters['L_0'].value = L_0_baseline


@earmbench
def fig_4b():
    print("Simulating model for figure 4B...")

    t = linspace(0, 6*3600, 6*60+1)  # 6 hours
    x = sim.run(tspan=t).all

    x_norm = c_[x['Bid_unbound'], x['PARP_unbound'], x['mSmac_unbound']]
    x_norm = 1 - x_norm / x_norm[0, :]  # gets away without max() since first values are largest

    # this is what I originally thought 4B was plotting. it's actually very close. -JLM
    #x_norm = array([x['tBid_total'], x['CPARP_total'], x['cSmac_total']]).T
    #x_norm /= x_norm.max(0)

    tp = t / 3600  # x axis as hours

    plt.figure("Figure 4B")
    plt.plot(tp, x_norm[:,0], 'b', label='IC substrate (tBid)')
    plt.plot(tp, x_norm[:,1], 'y', label='EC substrate (cPARP)')
    plt.plot(tp, x_norm[:,2], 'r', label='MOMP (cytosolic Smac)')
    plt.legend(loc='upper left', bbox_to_anchor=(0,1)).draw_frame(False)
    plt.xlabel('Time (hr)')
    plt.ylabel('fraction')
    a = plt.gca()
    a.set_ylim((-.05, 1.05))


# Run the code for each function 5 times
for _ in range(5):
    fig_4a()
    fig_4b()
```
