## Supplementary File S1 for "*Microbench*: Automated metadata management for systems biology benchmarking and reproducibility in Python": test_microbench_pysb.py

```
import tempfile
import os

def test_microbench_pysb():
    with tempfile.TemporaryDirectory() as td:
        # Set working directory to temp dir
        os.chdir(td)

        # Execute the script via import
        import microbench_pysb
```
