## Supplementary File S1 for "*Microbench*: Automated metadata management for systems biology benchmarking and reproducibility in Python": microbench-numpy.ipynb

```
# %% [markdown]
# Analyzing NumPy benchmarks using microbench

# %% [markdown]
## Background
This is a quick microbench example to demonstrate how two different versions of the NumPy library can give different outputs. The change highlighted here is a documented API change and not a bug, however, if a user runs code across heterogeneous compute environments (e.g. a cluster) it may not be immediately obvious that some nodes may be differently or misconfigured. A small Python program can be found in this directory, together with a JSON file containing microbench-captured metadata.

# %% [markdown]
## Load the data

# %% [code]
import pandas as pd

# %% [code]
results = pd.read_json('microbench-numpy.json', lines=True)

# %% [markdown]
## Show the captured metadata

# %% [code]
results

# %% [code]
results['runtime'] = results['finish_time'] - results['start_time']

# %% [code]
results['runtime']

# %% [markdown]
We can also reformat the `package_versions` dictionaries into separate columns, which is useful if we want to sort or filter by version number.

# %% [code]
results = pd.concat([results, pd.json_normalize(results['package_versions'])], axis=1).drop(columns='package_versions')

# %% [code]
results.head()

# %% [markdown]
## Discussion
The above example demonstrates different `return_value`s for NumPy 1.15 and 1.20. While environment standardization using virtual environments is encouraged, capturing metadata can help detect misconfigured environments.
```
