## Supplementary File S1 for "*Microbench*: Automated metadata management for systems biology benchmarking and reproducibility in Python": microbench_numpy.py

```
import numpy as np

# Microbench setup
from microbench import MicroBench, MBGlobalPackages, MBHostInfo, \
    MBReturnValue


class NumpyBench(MicroBench, MBGlobalPackages, MBHostInfo, MBReturnValue):
    outfile = 'microbench-numpy.json'


numpybench = NumpyBench()
# End microbench setup


@numpybench
def sum_linspace():
    """ Example with differing results by numpy version

    This code is based on an example given in the numpy v1.20.0 release notes:
    https://numpy.org/doc/stable/release/1.20.0-notes.html#np-linspace-on-integers-now-uses-floor

    As the numpy authors explain, "when using a int dtype in numpy.linspace,
    previously float values would be rounded towards zero. Now numpy.floor is
    used instead, which rounds toward -inf. This changes the results for
    negative values."

    On numpy <1.20, the returned value is -6. On numpy >=1.20, the returned
    value is -11.
    """
    return sum(np.linspace(-3, 1, 8, dtype=int))


print(sum_linspace())
```
