## Supplementary File S1 for "*Microbench*: Automated metadata management for systems biology benchmarking and reproducibility in Python": microbench-slurm.ipynb

```
# %% [markdown]
# Benchmarking Python code on SLURM using microbench

# %% [markdown]
## Background
SLURM is a cluster management system which handles queueing and resource allocation. Clusters are often extended over time and may contain heterogeneous hardware or software configuration, which may not be obvious to the end user. Here, we demonstrate microbench-captured hardware metadata (CPU frequency and RAM), and show that a demonstration Python script runs at vastly different speeds depending on the node CPU frequency (which may also correlate with other factors, e.g. later CPU generations, which we don't explore here). 

# %% [markdown]
## Load the data

# %% [code]
import pandas as pd
import seaborn as sb

# %% [code]
results = pd.read_json('microbench-slurm.json', lines=True)

# %% [markdown]
## Results

# %% [code]
# Calculate runtime for each run
results['runtime'] = results['finish_time'] - results['start_time']

# %% [code]
sbplot = sb.boxplot(x=results['cpu_freq_max'], y=results['runtime'].dt.total_seconds())

# %% [code]
sbplot = sb.boxplot(x=results['ram_total'], y=results['runtime'].dt.total_seconds())

# %% [markdown]
## Discussion
These results demonstrate how microbench can be used to analyze Python benchmarking data captured from a heterogeneous compute cluster, to identify how code performs on different hardware. Microbench makes the metadata capture and analysis process straightforward.
```
