## Supplementary File S1 for "*Microbench*: Automated metadata management for systems biology benchmarking and reproducibility in Python": microbench_slurm.py

```
#!/usr/bin/env python
"""
Python 3.6 script demonstrating vectorized execution
"""
import numpy as np
import time
import microbench

from microbench import *
import psutil


class MBHostCpuMaxFreq(microbench._NeedsPsUtil):
    """ Capture the number of logical CPU cores """
    def capture_cpu_cores(self, bm_data):
        self._check_psutil()
        bm_data['cpu_freq_max'] = psutil.cpu_freq().max

class MyBench(MicroBench, MBFunctionCall, MBPythonVersion, MBHostInfo, MBHostCpuMaxFreq, MBHostRamTotal):
    outfile = 'microbench_slurm.json'
    capture_versions = (np, )  # Or use MBGlobalPackages/MBInstalledPackages
    env_vars = ('SLURM_ARRAY_TASK_ID', )
    
mybench = MyBench()


@mybench
def accre_demo():
    # 10 million entries
    t = np.linspace(-10,10,10000000)
    x1 = np.zeros(len(t))
    x2 = np.zeros(len(t))

    time1 = time.process_time()
    # naive, non-vectorized implementation
    for i, ti in enumerate(t):
        x1[i] = np.sin(ti)
    time2 = time.process_time()
    print(f"naive implementation {time2-time1:.2f} cpu seconds elapsed")

    # vectorized implementation
    time1 = time.process_time()
    x2 = np.sin(t)
    time2 = time.process_time()
    print(f"vectorized implementation {time2-time1:.2f} cpu seconds elapsed")

    if np.array_equal(x1,x2):
        print("arrays equal!")

accre_demo()
```
