## Supplementary File S1 for "*Microbench*: Automated metadata management for systems biology benchmarking and reproducibility in Python": microbench_slurm.sh

```
#!/bin/bash
#SBATCH --ntasks=1
#SBATCH --time=00:10:00
#SBATCH --mem=256M
#SBATCH --array=1-100
#SBATCH --output=microbench_slurm.out

# Create the virtual environment first using the separate create-venv.sh script
source mbenv/bin/activate

python microbench_slurm.py
```
