## Supplementary File S1 for "*Microbench*: Automated metadata management for systems biology benchmarking and reproducibility in Python": README.md

# Microbench examples

[![Binder](https://mybinder.org/badge_logo.svg)](https://mybinder.org/v2/gh/alubbock/microbench-examples/HEAD)

Example use cases for the metadata capture, benchmarking, and reproducibility
package [microbench](https://github.com/alubbock/microbench).
Microbench captures runtime metadata from a Python function, such as package
versions and timing information, which are saved to a JSON file or a Redis
instance.

Each example contains a Jupyter Notebook which shows how to analyze the JSON
file and perform some basic visualizations. Within each example is a `src`
directory which contains the original Python program used to generate the
metadata.

The three examples are:

* A [PySB](https://pysb.org) model called [EARM](https://earm.readthedocs.io)
* A NumPy example showing a (documented) reproducibility difference across
  versions due to a
  [NumPy API change](https://numpy.org/doc/stable/release/1.20.0-notes.html#np-linspace-on-integers-now-uses-floor)
* A [SLURM](https://slurm.schedmd.com) example showing how metadata can be
  caputed for jobs run on a cluster/scheduling system
